# Conjugative transfer of naturally occurring plasmids in *Mycolicibacterium* sp

**DOI:** 10.1101/2021.10.22.464858

**Authors:** Sergio Morgado, Ana Carolina Vicente

## Abstract

Conjugation is considered the main horizontal gene transfer (HGT) mechanism in bacterial adaptation and evolution. In the *Mycobacteriaceae* family, *Mycolicibacterium smegmatis* has been used as the model organism for the conjugative transfer of hybrid plasmids. However, the natural conjugation process in any bacteria would involve the transfer of naturally occurring plasmids. Currently, there is a gap in this regard in relation to this abundant environmental genus of *Mycobacteriaceae*. Here, we performed conjugation experiments between wild *Mycolicibacterium* sp. strains involving naturally occurring plasmids (sizes of 21 and 274 kb), and interestingly, evidence of conjugative transfer was obtained. Thus, it is likely that conjugation occurs in *Mycolicibacterium* in the natural environment, representing a source of diversification and evolution in this genus of bacteria.

## Introduction

Horizontal gene transfer is fundamental in bacterial adaptation and evolution, occurring through three natural processes: conjugation, transduction, and transformation. Among them, conjugation occurs through direct cell-cell contact, and it is the main mechanism that contributes to the plasmid dispersion, as it occurs between several bacterial phyla (being mainly studied in *Proteobacteria*) (Wang et al., 2003; Kohler et al., 2019). In the *Mycobacteriaceae* family, which includes the *Mycobacterium* genus and four recently reclassified genera (*Mycolicibacterium, Mycobacteroides, Mycolicibacter*, and *Mycolicibacillus*) (Gupta et al., 2018), conjugation has already been associated with plasmids and chromosomal fragments (distributive conjugal transfer) (Gray and Derbyshire, 2018). The mycobacterial conjugation system is driven by the type VII secretion system (T7SS), which is encoded by six paralogous loci (ESX-1, -2, -3, -4, -5 and -4-bis), each one with different genetic organizations and functions (Dumas et al., 2016). Due to the impact that plasmid exchange has on many aspects of bacterial biology, this issue has been studied in depth in several bacterial taxa, but in *Mycobacteriaceae* little significance has been given to this point (Shoulah et al., 2018).

Plasmid conjugation in *Mycobacteriaceae* has only been experimentally observed in few species of three genera: *Mycobacterium* (*M. tuberculosis, M. marinum, M. avium, M. kansasii*, and *M. bovis*), *Mycobacteroides* (*M. abscessus*), and *Mycolicibacterium* (*M. smegmatis*). Moreover, conjugations in these genera only occurred between organisms of the same genus, species, or to *Escherichia coli* (Wang et al., 2003; Rabello et al., 2012; Leão et al., 2013; Ummels et al., 2014; Gray and Derbyshire, 2018; Shoulah et al., 2018). Concerning *Mycolicibacterium* genus and plasmid conjugation, only one species, *M. smegmatis*, has been considered as a model in tests with recombinant plasmids (lacks natural plasmids), since it is a fast-growing and non-pathogenic species (Lazraq et al., 1990; Wang et al., 2003; Derbyshire and Gray, 2014; Gray and Derbyshire, 2018). In this genus, plasmids were thought to be scarce (Gray and Derbyshire, 2018; Morgado and Vicente, 2021) and reports of conjugative transfer of naturally occurring plasmids are rare, only showing successful conjugation of a small (<10 kb size) non-conjugative plasmid from *E. coli* to *M. smegmatis* (Gormley and Davies, 1991).

Previous genomic analyzes on *Mycolicibacterium* sp. from Atlantic Forest soil revealed the presence of three plasmids (pCBMA213_1, ∼274 kb; pCBMA213_2, ∼160 kb; and pCBMA213_3, ∼21 kb) in a lineage. Curiously, strains from this lineage presented distinct plasmid profiles, varying from one (pCBMA213_3) to the three plasmids. In addition, in another *Mycolicibacterium* sp. lineage, no plasmids were identified (Morgado and Vicente, 2020). So, to contribute with experimental evidence of conjugation in this bacterial family, analyzing the conjugative capability of these natural plasmids, we performed conjugation tests using these wild *Mycolicibacterium* strains carrying plasmids as donors and a wild *Mycolicibacterium* as the recipient. In this way, we revealed evidence of conjugative transfer of naturally occurring plasmids in the *Mycolicibacterium* genus.

## Materials and Methods

### Bacterial strains

Four *Mycolicibacterium* sp. strains (*Mycolicibacterium* sp. CBMA213, *Mycolicibacterium* sp. CBMA234, *Mycolicibacterium* sp. CBMA311, and *Mycolicibacterium* sp. CBMA360) isolated from Atlantic Forest soil (deposited in CBAS, Bacterial Collection, Fiocruz/Brazil) were used in this study. They were cultivated in 5 ml of Tryptic Soy Broth (TSB) supplemented with Tween80 (0.05%) for 10 days at 22°C before the mating experiments.

### Pulsed-field gel electrophoresis

Genomic DNA of the *Mycolicibacterium* strains was submitted to PFGE. After bacterial growth, cells were suspended in PIV buffer (10 mM Tris-HCl pH 7.6, 1 M NaCl). Agarose plugs were prepared by mixing the suspension in PFGE molds, and after solidification, the plugs were transferred to lysis buffer (6 mM Tris-HCl pH 7.6, 1 M NaCl, 100 mM EDTA, 0.2% deoxycholate, 0.5% N-lauroylsarcosine, 0.5% Brij-58, 10 mg/ml lysozyme) and incubated overnight at 37°C. Lysis buffer was removed and the plugs were incubated overnight at 50°C in ESP buffer (0.5 M EDTA pH 8, 1% N-lauroylsarcosine, 100 µg/ml proteinase K). Next, the plugs were washed four times/day for a week with TE buffer (10 mM Tris-HCl pH 8, 0.1 mM EDTA) until used. The plugs were loaded on a 1.2% agarose gel in a Bio-Rad CHEF-DR III system containing TBE buffer (44.5 mM Tris-HCl, 44.5 mM boric acid, 1 mM EDTA, pH 8.3), and were subjected to a pulse of 5 s ramping to 35 s for 18 h at 5.5 V/cm. Two runs were carried out: one with non-digested DNA, and the other with digested DNA by DraI restriction enzyme (Promega) at 37°C overnight. In the gels, the Lambda ladder PFG marker (New England BioLabs) was used as molecular standard.

### Mating experiments

Mating experiments were performed in triplicates using *M*. sp. CBMA213, *M*. sp. CBMA311, and *M*. sp. CBMA360 as donors, and *M*. sp. CBMA234 as the recipient. The recipient strain was negative for the presence of plasmids, as observed by PCR of plasmid genes and non-digested PFGE. Aliquots of 2 ml of each recipient and donor strains were combined (three mating pairs in total) and filtered through sterile 0.22 µm membranes (Merck Millipore, GSWP02500), which were incubated over TSA plates for 8 days at 22°C. After this period, the membranes were transferred to sterile containers and washed with 5 ml of TSB supplemented with 5 µg/ml of rifampicin (Sigma-Aldrich). Next, serial dilutions were plated on TSA supplemented with 5 µg/ml of rifampicin. Rifampicin was used as a selection marker because the donor cells were susceptible on 5 µg/ml concentration, while the recipient, not. The putative transconjugants were pooled from the TSB plates and submitted to further analysis. Those pools that did not generate amplicons related to the gene markers of the donor strains and presented a PFGE-DraI profile equal to the recipient strain were assumed to be transconjugants. The presence of the plasmids in transconjugants was verified by non-digested PFGE and amplification of plasmid gene markers.

### PCR assays

PCR assays were performed using different PCR kits, depending on the set of primers used (Table 1), in 50 µl reactions in the following conditions: (i) 95°C for 5 min, 40 cycles of 95°C for 30 s, 55°C for 30 s and 72°C for 45 s, with a final elongation step at 72° for 10 min (Promega, GoTaq DNA Polymerase); (ii) 98°C for 5 min, 40 cycles of 96°C for 1 min, 50°C for 30 s and 72°C for 40 s, with a final elongation step at 72° for 10 min (Qiagen, Taq DNA Polymerase). The amplicons were purified using GFX PCR DNA and Gel Band Purification Kit (Sigma-Aldrich), according to the manufacturer, and the products were sequenced by 3730XL DNA Analyser (Applied Biosystems).

**Table 1.** Primer sequences

| Primers | Sequences (5' -> 3') | Gene target | Amplicon size (bp) | PCR kit |
| --- | --- | --- | --- | --- |
| cbma234U | agc atc gct gag ttc aag g (F)<br>tta gct gtt tga ccc tgc tg (R) | *arr* | 630 | Promega |
| cbma213U | gac cgg acc tga atg ttc tt (F)<br>gtg cgt tat caa tcg tcc tc (R) | hypothetical | 606 | Promega |
| pCBMA213_1 | gcg atg agg aac ggt act aaa (F)<br>cgc tcc ata gtt gtc atg ct (R) | *vap* | 526 | Promega |
| pCBMA213_2 | atc tcc cgt aag acg ctg at (F)<br>ccg ggg gta gtt gtt tta tc (R) | *rep* | 338 | Qiagen |
| pCBMA213_3 | gca atg tgg tga tcc tga gt (F)<br>aca aga agg gca tga gca (R) | hypothetical | 332 | Promega |

## Results

The experiments were performed using three donor strains, which presented different plasmid profiles, as observed in previous *in silico* analyzes (Morgado and Vicente, 2020): *Mycolicibacterium* sp. CBMA213; bearing pCBMA213_1, pCBMA213_2, and pCBMA213_3; *Mycolicibacterium* sp. CBMA311, bearing pCBMA213_3; and *Mycolicibacterium* sp. CBMA360, bearing pCBMA213_2 and pCBMA213_3. The recipient strain was *Mycolicibacterium* sp. CBMA234, which is devoid of plasmids. To validate these *in silico* plasmid predictions, the plasmid profiles of the strains were revealed by pulsed-field gel electrophoresis of non-digested DNA, which showed the presence of different band profiles (plasmids) in the donor strains, and the absence of bands in the recipient strain (Figure 1).

**Figure 1.**
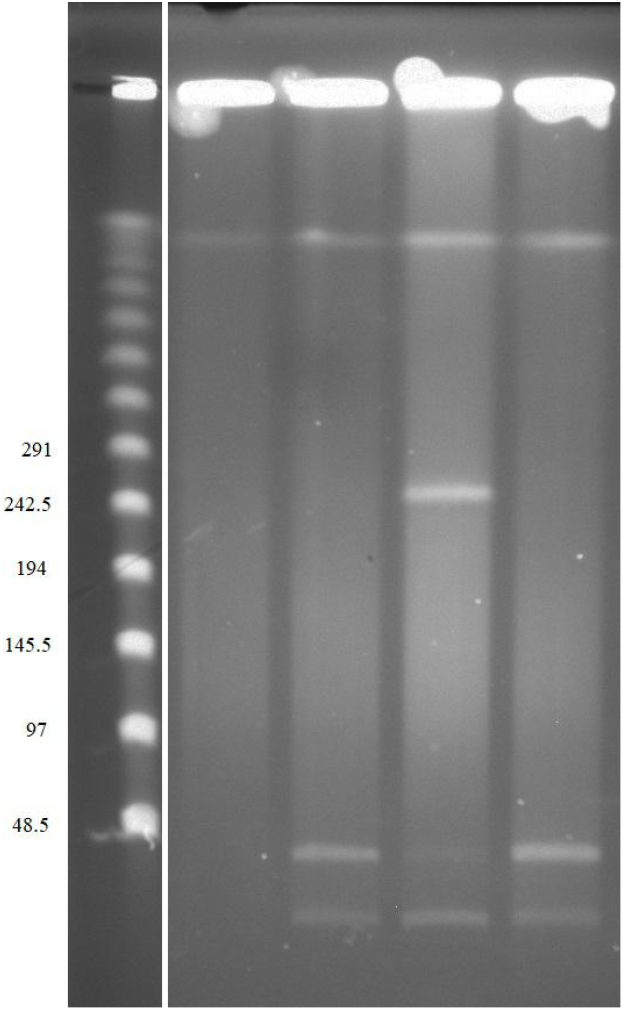
PFGE with undigested DNA from *M*. sp. CBMA234, *M*. sp. CBMA311, *M*. sp. CBMA213, *M*. sp. CBMA360. In the left column the size (kb) of the bands is shown.

In addition to PFGE, we also showed the presence of the distinct plasmids in the donor strains by PCR targeting marker genes of each plasmid (Table 2). These marker genes were assigned to these plasmids based on the *in-silico* analyses (Morgado and Vicente, 2020). The plasmid pCBMA213_2 (∼160 kb size) was not observed in the PFGE, however, the presence of this plasmid, and the others, in the donor strains was previously characterized by whole genome sequencing (Morgado and Vicente, 2020), and here by PCR (Table 2).

**Table 2.**
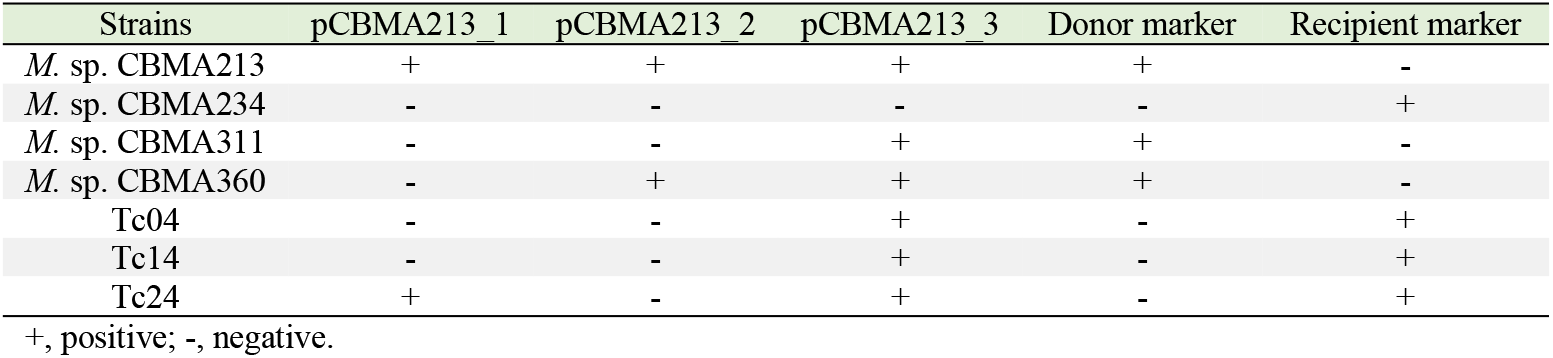
PCR results for the marker genes of each strain

| Strains | pCBMA213_1 | pCBMA213_2 | pCBMA213_3 | Donor marker | Recipient marker |
| --- | --- | --- | --- | --- | --- |
| *M. sp.* CBMA213 | + | + | + | + | - |
| *M. sp.* CBMA234 | - | - | - | - | + |
| *M. sp.* CBMA311 | - | - | + | + | - |
| *M. sp.* CBMA360 | - | + | + | + | - |
| Tc04 | - | - | + | - | + |
| Tc14 | - | - | + | - | + |
| Tc24 | + | - | + | - | + |

+, positive; -, negative.

The mating experiments were performed on a solid medium using *M*. sp. CBMA311, *M*. sp. CBMA360, or *M*. sp. CBMA213 as donors, and *M*. sp. CBMA234 as the same recipient. The putative transconjugants were selected in a solid medium supplemented with 5 µg/ml of rifampicin. The plasmids pCBMA213_1, pCBMA213_2, and pCBMA213_3 lack a suitable marker for the selection of eventual transconjugants. However, the recipient and donor strains could be selected by their rifampicin susceptibility, since the recipient strain is resistant to ≥ 5 µg/ml of rifampicin, while the donors do not grow at this concentration (data not shown). To check the success of the conjugation, the transconjugant pools (Tc04, Tc14, and Tc24: transconjugants of the recipient *M*. sp. CBMA234 with the donors *M*. sp. CBMA360, *M*. sp. CBMA311, or *M*. sp. CBMA213, respectively) were submitted to PFGE of non-digested DNA and PCR. As PFGE result, no bands corresponding to the plasmids were revealed. Conversely, a PCR assay, based on the plasmid marker genes, resulted in amplicons of pCBMA213_3 plasmid (∼21 kb) in Tc04, Tc14, and Tc24 pools; and pCBMA213_1 plasmid (∼274 kb) in Tc24 pool (Table 2). These amplicons were further Sanger sequenced and confirmed as the target genes. In addition, to verify the absence of the donor strain in the transconjugant pools, a PFGE with DraI restriction enzyme digestion was performed. The wild donors and the recipient, as well as transconjugants, had their PFGE pattern defined, and it was possible to observe that the transconjugants share the same pattern with the recipient strain (Figure 2). Furthermore, the transconjugant pools were submitted to PCR targeting marker genes of the donor and recipient strains, revealing the presence of only the recipient strain (Table 2). Altogether, these results showed that pCBMA213_1 and pCBMA213_3 were transferred by a conjugation-like mechanism between *Mycolicibacterium* strains.

**Figure 2.**
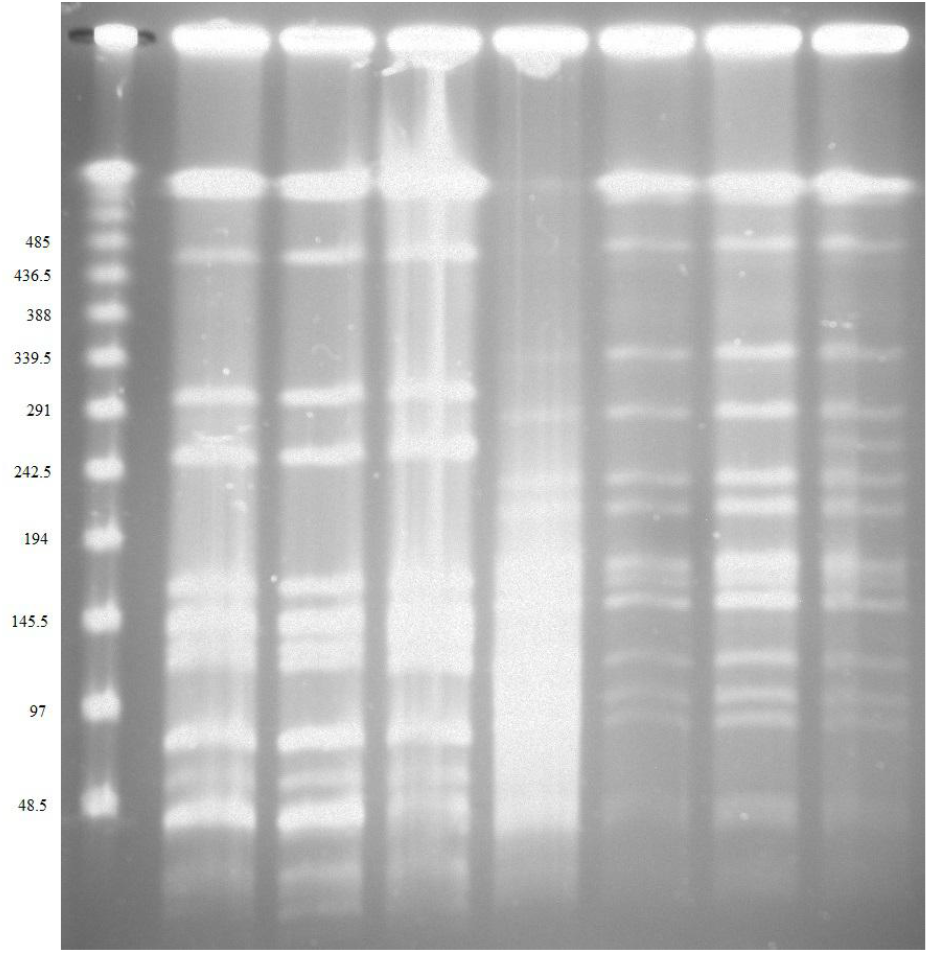
PFGE with DraI digested DNA from *M*. sp. CBMA360, *M*. sp. CBMA311, *M*. sp. CBMA213, *M*. sp. CBMA234, Tc04, Tc14, and Tc24. In the left column the size (kb) of the bands is shown.

## Discussion

To date, the experimental studies of HGT in *the Mycobacteriaceae* family involved only organisms of few species. Among them, *Mycolicibacterium smegmatis* is the model organism used in several conjugative experiments involving transfer of recombinant (successful) and natural (unsuccessful) plasmids to/from other genera of *Mycobacteriaceae* (Lazraq et al., 1990; Gormley and Davies, 1991; Wang et al., 2003; Rabello et al., 2012; Leão et al., 2013; Ummels et al., 2014; Derbyshire and Gray, 2014). Here, we present experimental evidence of conjugation of naturally occurring plasmids between wild *Mycolicibacterium* sp. strains, both as a recipient and as a donor. Among the three plasmids tested, two of them (pCBMA213_1 and pCBMA213_3) could be observed in the transconjugant strains, showing that *Mycolicibacterium* can transfer plasmids with a wide variation in size (21 - 274 kb) in a conjugation-like mechanism. In fact, the transfer efficiency appears to have been low, as the transconjugants were only detected through molecular assays. In previous studies, in which donors and recipients were *Mycobacterium* strains, it was demonstrated conjugation-like mechanisms with low and high transfer efficiency (Rabello et al., 2012; Ummels et al., 2014; Shoulah et al., 2018). However, conjugation experiments between *Mycolicibacterium* and other *Mycobacteriaceae* genera (*Mycobacterium* or *Mycobacteroides*) were not successful (Rabello et al., 2012; Leão et al., 2013; Ummels et al., 2014), suggesting the existence of some species barrier (Neil et al., 2021).

In a previous genomic analysis of the two successfully transferred plasmids (Morgado and Vicente, 2020), no classical conjugative gene was identified, such as *vir*B4, *vir*D4, and relaxases (Smillie et al., 2010). Moreover, sequences associated with known plasmid origin of transfer (oriT) were not identified (Morgado and Vicente, 2020), which would define them as non-mobilizable plasmids (Smillie et al., 2010). In *the Mycobacterium* genus, a conjugative plasmid was shown to require both type IV and type VII secretion systems (ESX-2) (Ummels et al., 2014). Curiously, pCBMA213_1 harbors a distinct T7SS (ESX-3), but it lacks type IV secretion system genes (Morgado and Vicente, 2020). Although pCBMA213_1 and pCBMA213_3 plasmids were characterized as non-mobilizable plasmids, they were transferred, so it is likely that they were transferred *in trans* by other elements through unknown oriT sites or that they have an as-yet-undescribed conjugation mechanism. Thus, pCBMA213_2 could act as a helper plasmid, mobilizing other plasmids (Guédon et al., 2017), since this plasmid carries the set of genes associated with conjugation (Morgado and Vicente, 2020). Indeed, the *E. coli* plasmids transferred to *M. smegmatis* were supported by helper plasmids (Lazraq et al., 1990; Gormley and Davies, 1991). Another alternative could involve the distributive conjugal transfer mechanism, which involves the ESX-1 and ESX-4 secretion systems, and it has been associated with mycobacterial conjugation of unlinked chromosomal fragments (Wang et al., 2003; Gray and Derbyshire, 2018). Interestingly, here, the chromosome of the donor strains carried ESX-4, while the chromosome of the recipient strain had both ESX-1 and ESX-4.

In conclusion, these findings provide new evidence for conjugative transfer of naturally occurring plasmids in the *Mycolicibacterium* genus, as other transfer mechanisms, such as transformation, seem unlikely to have occurred due to the sizes of transferred plasmids. These evidence are clues that can be further explored in this diverse and important family of bacteria.

## Acknowledgments

This work was supported by the Coordination for the Improvement of Higher Education Personnel (Coordenação de Aperfeiçoamento de Pessoal de Nível Superior—CAPES)—Finance Code 001; and Inova Fiocruz/Fundação Oswaldo Cruz.

## References

Wang J, Parsons LM, Derbyshire KM. Unconventional conjugal DNA transfer in mycobacteria. Nat Genet 2003, doi:10.1038/ng1139.

Kohler V, Keller W, Grohmann E. Regulation of Gram-Positive Conjugation. Front Microbiol 2019, doi:10.3389/fmicb.2019.01134.

Gupta RS, Lo B, Son J. Phylogenomics and Comparative Genomic Studies Robustly Support Division of the Genus Mycobacterium into an Emended Genus Mycobacterium and Four Novel Genera. Front Microbiol 2018, doi:10.3389/fmicb.2018.00067.

Gray TA, Derbyshire KM. Blending genomes: distributive conjugal transfer in mycobacteria, a sexier form of HGT. Mol Microbiol 2018, doi:10.1111/mmi.13971.

Dumas E, Christina Boritsch E, Vandenbogaert M et al. Mycobacterial Pan-Genome Analysis Suggests Important Role of Plasmids in the Radiation of Type VII Secretion Systems. Genome Biol Evol 2016, doi:10.1093/gbe/evw001.

Shoulah SA, Oschmann AM, Selim A et al. Environmental Mycobacterium avium subsp. hominissuis have a higher probability to act as a recipient in conjugation than clinical strains. Plasmid 2018, doi:10.1016/j.plasmid.2018.01.003.

Rabello MC, Matsumoto CK, Almeida LG et al. First description of natural and experimental conjugation between Mycobacteria mediated by a linear plasmid. PLoS One 2012, doi:10.1371/journal.pone.0029884.

Leão SC, Matsumoto CK, Carneiro A et al. The detection and sequencing of a broad-host-range conjugative IncP-1β plasmid in an epidemic strain of Mycobacterium abscessus subsp. Bolletii. PLoS One 2013, doi:10.1371/journal.pone.0060746.

Ummels R, Abdallah AM, Kuiper V et al. Identification of a novel conjugative plasmid in mycobacteria that requires both type IV and type VII secretion. mBio 2014, doi:10.1128/mBio.01744-14.

Lazraq R, Clavel-Sérès S, David HL et al. Conjugative transfer of a shuttle plasmid from Escherichia coli to Mycobacterium smegmatis. FEMS Microbiol Lett 1990, doi:10.1016/0378-1097(90)90427-r.

Derbyshire KM, Gray TA. Distributive Conjugal Transfer: New Insights into Horizontal Gene Transfer and Genetic Exchange in Mycobacteria. Microbiol Spectr 2014, doi:10.1128/microbiolspec.MGM2-0022-2013.

Morgado SM, Vicente ACP. Comprehensive in silico survey of the Mycolicibacterium mobilome reveals an as yet underexplored diversity. Microb Genom 2021, doi:10.1099/mgen.0.000533.

Gormley EP, Davies J. Transfer of plasmid RSF1010 by conjugation from Escherichia coli to Streptomyces lividans and Mycobacterium smegmatis. J Bacteriol 1991, doi:10.1128/jb.173.21.6705-6708.1991.

Morgado SM, Paulo Vicente AC. Genomics of Atlantic Forest Mycobacteriaceae strains unravels a mobilome diversity with a novel integrative conjugative element and plasmids harbouring T7SS. Microb Genom 2020, doi:10.1099/mgen.0.000382.

Neil K, Allard N, Rodrigue S. Molecular Mechanisms Influencing Bacterial Conjugation in the Intestinal Microbiota. Front Microbiol 2021, doi:10.3389/fmicb.2021.673260.

Smillie C, Garcillán-Barcia MP, Francia MV et al. Mobility of plasmids. Microbiol Mol Biol 2010, doi:10.1128/MMBR.00020-10.

Guédon G, Libante V, Coluzzi C et al. The Obscure World of Integrative and Mobilizable Elements, Highly Widespread Elements that Pirate Bacterial Conjugative Systems. Genes (Basel) 2017, doi:10.3390/genes8110337.

